# Neurophysiological correlates of cortical hierarchy across the lifespan

**DOI:** 10.1101/2024.07.15.602693

**Authors:** Jana Fehring, Elio Balestrieri, Niels K. Focke, Udo Dannlowski, Christina Stier, Joachim Gross

**Author notes:** joined senior authors (Christina Stier, Joachim Gross).

## Abstract

The brain processes information along a hierarchical structure, forming a gradient of cortical hierarchy from sensorimotor areas to transmodal areas. Here, we aim to understand which aspects of neural dynamics characterize this gradient and whether the respective spatial distribution varies across the lifespan. Therefore, we extracted neurophysiological features from magnetoencephalography recordings in 350 participants between 18 and 88 years during rest. Among traditional features related to the power spectrum, delta power (1-4 Hz) showed the most robust association with cortical hierarchy, increasing along this axis. Beyond traditional features, we employed comprehensive time-series characterization and identified a novel hierarchy-sensitive feature capturing the variability of the signal’s mean over time. This feature increases along the cortical hierarchy, suggesting that higher-level brain areas exhibit more dynamic and context-dependent activity patterns. Furthermore, we highlight changes in the gradient of brain dynamics across the lifespan. Alpha power distribution, for instance, exhibits a posterior-anterior gradient in young adults that becomes less pronounced with increasing age. Further, the change of the autocorrelation and auto mutual information function along the cortical hierarchy is heavily modulated by age. These findings reveal simple but robust neurophysiological markers of cortical hierarchy and highlight the dynamic nature of the brain’s organization throughout life.

## Introduction

The brain processes incoming stimuli, e.g. visual stimuli, with increasingly complex representations of the original stimulus along the visual processing pathways. While more primary brain areas represent simpler properties of the stimulus, the representation of the stimulus becomes increasingly abstract in higher-order brain areas where it also is integrated with stimuli from other modalities ^1–5^. This processing hierarchy is spatially distributed across the cortex forming an axis of cortical hierarchy. This axis is characterized by gradual changes in various anatomical and functional measures from primary sensorimotor regions (encompassing somatosensory, motor, auditory, and visual areas) to transmodal brain areas, including the prefrontal, temporal, as well as inferior parietal cortex ^6–8^. Here, primary brain areas typically occupy spatially distal positions, with transmodal regions located between them ^7,9^. Biological measures forming the brain’s structure that change gradually and consistently along the hierarchy include myelination ^10–12^, cortical thickness ^11,13^, gene expression ^11,14–16^, neurotransmitter receptor densities ^17–19^, neuron density ^20,21^ as well as neuron spine density ^22^. As the brain’s structure is both the scaffold and constraint of the function of the brain, differentiation along the cortical hierarchy can be also observed in measures of brain activity. For instance, the first principal gradient of functional connectivity, measured using fMRI, spans from unimodal to transmodal areas ^9^. Moreover neurophysiological measures, such as peak frequency, the 1/f exponent of the aperiodic decay ^23^, and the timescales of intrinsic fluctuations ^22,24–29^ reflect systematic changes in regional neurophysiological signals that are likely driven by the underlying anatomical changes, in turn supporting the specialized functions of brain areas across the hierarchy.

So far, only few studies have examined the electrophysiological correlates of neural dynamics at different hierarchical levels ^22,24–29^ and used a restricted set of features for characterizing brain activity. This trend has been only recently reversed in a work by Shafiei et al. ^30^: In this work, MEG-derived time series were deeply phenotyped. This approach disclosed that linear combinations of time-series features significantly correlate with linear combinations of diverse micro-architectural features such as neurotransmitters, gene expression, T1w/T2w ratio, and others ^30^. Nonetheless a clear understanding of which aspects of neural dynamics characterize cortical hierarchy is still missing.

Importantly, critical differences in the brain exist not only spatially, but also in time. Across the lifespan, both anatomical features such as gray and white matter volume ^31^, cortical thickness ^31–35^, myelination ^36^, spatial receptor distribution ^37–39^, as well as neural dynamics ^40–51^ are known to undergo important changes. Similarly, age-related changes have been described in the spatial distribution of the first principal gradient of functional connectivity, as measured with fMRI. A functional gradient describing cortical hierarchy emerges only during development ^52,53^ and disperses during aging ^54^. However, it remains to be determined which specific characteristics in the neural dynamics change with age in relation to the spatial gradient of cortical hierarchy.

To address these questions, we capitalized on traditional features used to describe MEG-derived time-series, such as spectral power, peak frequency, and 1/f exponent, as well as a recently developed approach for comprehensive time-series phenotyping ^55,56^. This allows the characterization of time-series by a set of several thousands of features that quantify the distribution, stationarity, correlation, complexity, non-linearity, predictability, and many more ^55,56^. Specifically, we characterized neural dynamics of source-localized MEG signals in atlas-defined brain areas. Here, we further build on previous work ^30^ to specifically study the relationship between cortical hierarchy and single time-series features in an age-stratified sample of 350 participants, and how this relationship changes with age. Our analysis reveals that simple time-series features align significantly with hierarchical level and capture age-related changes in cortical hierarchies.

## Results

To investigate which aspects of regional neural dynamics consistently follow the gradient of cortical hierarchy, we extracted and characterized traditional features related to the power spectrum as well as 5961 additional time-series features leveraging the hctsa toolbox ^55,56^ (Figure 1). These were computed on five minutes of source-reconstructed MEG resting-state data (eyes closed) in 200 parcels of the Schaefer atlas of 350 participants with an age range between 18 and 88 years from the CamCAN repository (Figure 1). For defining cortical hierarchy two different approaches were used. First, we used one archetypal map according to Sydnor et al. ^8^ integrating ten different measures related to cortical hierarchy, from hereon named cortical hierarchy for simplicity. Second, we used participant-specific cortical thickness data as a proxy for cortical hierarchy. We also employed two statistical approaches to test for significant associations between each of these two hierarchy maps and the individual time-series features. Linear-mixed effect models (LMEMs) allowed us to model interindividual differences and to include individual age in the model. The LMEM analysis was based on the following model: *time-series feature ∼ cortical hierarchy + age + cortical hierarchy*age + (1|subj).* Meanwhile, the second statistical approach, spin testing, specifically accounts for the typical spatial auto-correlation structure of brain maps. In this test, the original feature map is compared to a large number of surrogate maps that preserve spatial autocorrelation ^57,58^ (Figure 1). In the following, we report both of these complementary statistics.

**Figure 1:**
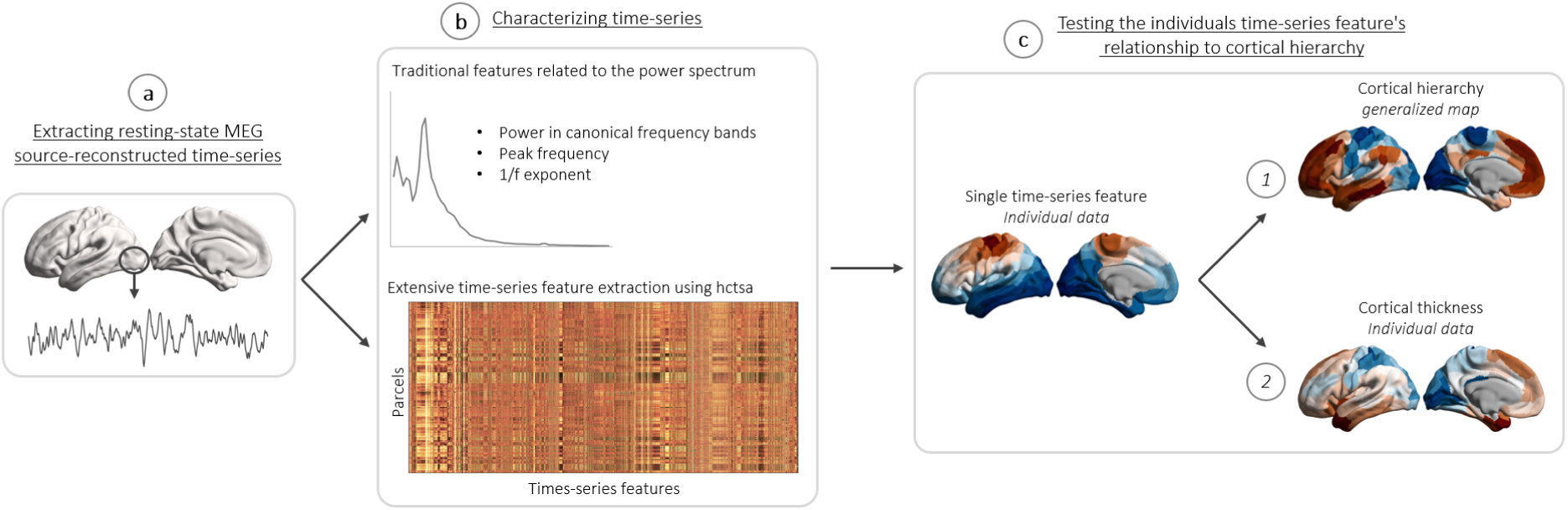
Schematic outline of the methodological approach. a: extracting MEG resting-state source-reconstructed time-series. For each participant, five minutes of MEG resting-state recording, eyes closed, have been source-reconstructed to 200 parcels according to the Schaefer atlas. b: the time-series of each parcel was characterized using traditional features which are related to the power spectrum (spectral power in the canonical frequency bands, alpha peak frequency, and 1/f exponent) as well as using extensive feature extraction leveraging the hctsa toolbox ^55,56^. The latter yielded 5961 additional time-series-related features. Each measure was computed on the parcel level for each participant, here displayed for one participant. c: Testing the association of the distribution of each feature across the cortex with cortical hierarchy. For this purpose, two definitions of cortical hierarchy were employed. 1: a generalized, archetypal map of cortical hierarchy incorporating multiple hierarchy-related measures and 2: participant-specific cortical thickness data.

### Low frequency spectral power is an index of cortical hierarchy

The analysis of resting-state neurophysiological data often relies on spectral power in canonical frequency bands. Therefore, we first set out to test to what extent frequency-specific power varies systematically with cortical hierarchy. We complemented these power features with two more advanced features that have recently gained interest in the literature, namely the slope of the power spectrum (1/f exponent) ^59^, and the frequency showing maximum power in the alpha band (8-13 Hz), namely alpha peak frequency. Thus, we computed spectral power in the canonical frequency bands (delta, theta, alpha, beta and gamma), as well as 1/f exponent, and peak frequency for each participant and cortical parcel. Subsequently, LMEMs and spin tests were computed for each feature using first, the archetypal map of cortical hierarchy and second, individual cortical thickness data (Figure 1c).

Inspection of brain maps and statistical results in Figure 2 revealed that beta power, 1/f exponent and peak frequency did not show a systematic relationship to cortical hierarchy. For the remaining features, both positive and negative correlations and LMEM t-values were observed. Delta, theta and gamma power increased with cortical hierarchy whereas alpha power decreased. These four frequency bands revealed significant associations with cortical hierarchy in the LMEM model. However, only delta and theta power showed significant FDR-corrected results for both hierarchy maps. Across all tests, the delta band emerged as the frequency band in which power has the strongest relationship with both the archetypal cortical hierarchy topography and the individual cortical thickness data (for full LMEM results, see Supplementary Table S2, S3). Increased delta power is found especially in prefrontal as well as temporal brain areas while especially in motor and visual areas delta power is low. T-values of the LMEM were generally very high and likely inflated by spatial autocorrelations of feature maps. We therefore computed 50000 surrogate feature maps that preserved spatial autocorrelation and recomputed LMEMs. The delta map is still significantly related to cortical hierarchy after accounting for spatial autocorrelation (Supplementary Figure S1a).

**Figure 2:**
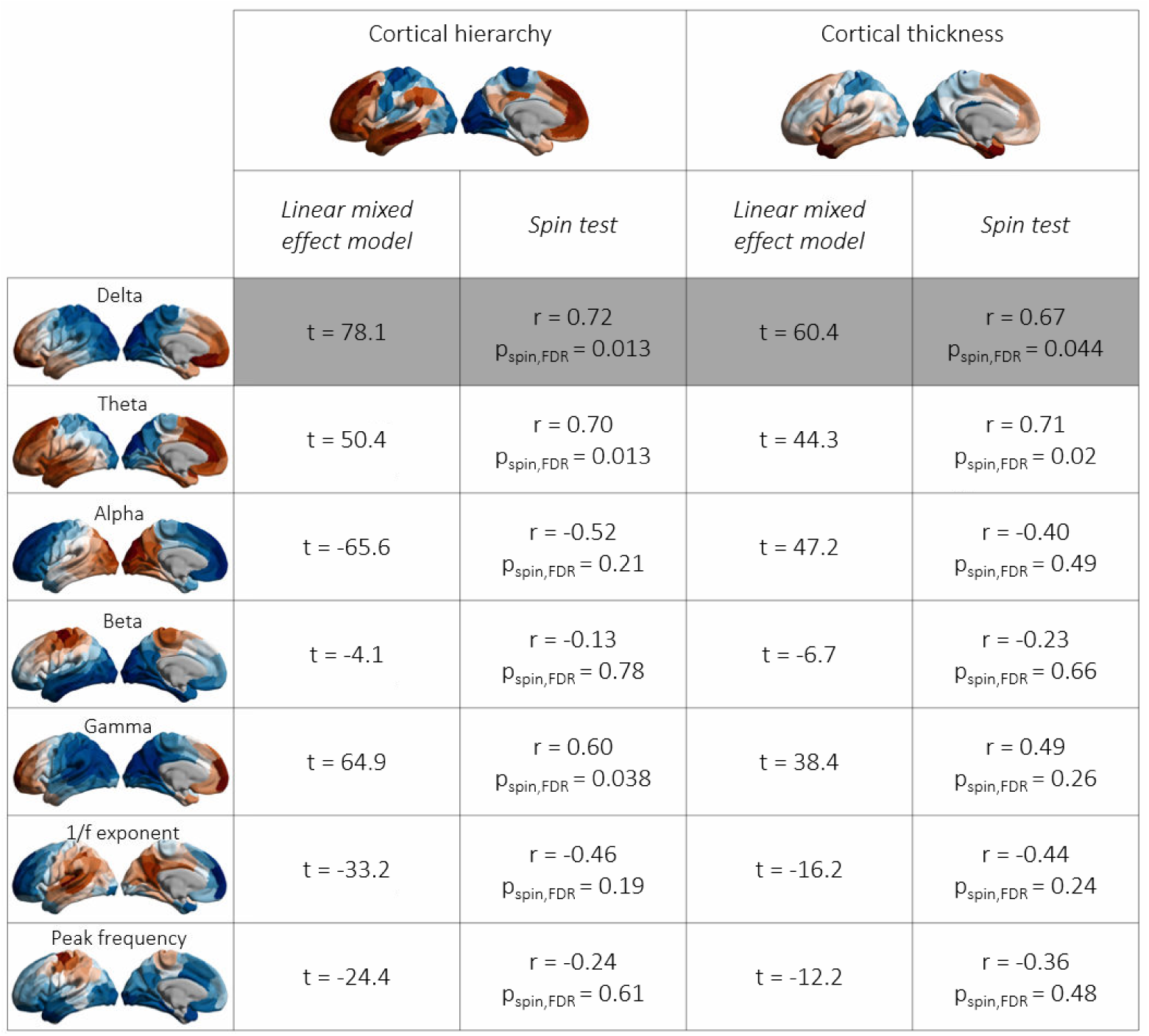
Statistical results for the relationship of traditional time-series features to cortical hierarchy. Each time-series feature (rows) was tested against a generalized, archetypal, topography of cortical hierarchy as well as cortical thickness, a proxy for cortical hierarchy. For each association, two different statistical approaches were employed, namely linear-mixed effect modeling and spin testing. For the linear mixed-effect model, here, only the main effect of cortical hierarchy or cortical thickness was listed. The p-values resulting from the spin testing were FDR corrected. Overall, the delta frequency band showed the most robust statistical effects across all performed tests, therefore a close relationship to cortical hierarchy. Colorscale: each time-series feature map is a group-averaged z-scored feature. Parcels with the highest feature value are colored red, while cortical parcels with low values are colored in blue.

### The variability of the mean increases along cortical hierarchy

In addition to traditional time-series features related to the power spectrum, massive feature extraction was performed in which 5961 additional time-series features were computed. Many of these features vary significantly along the axis of the archetypal cortical hierarchy map, indicated by high t-statistics in the LMEM. In order to avoid describing features which are displaying similar neural mechanisms, we performed pairwise correlation analysis and cluster analysis with the 50 features showing the highest t-values (Figure 3a; for more details, see Supplementary Figure S2, Table S1). Many of these features are highly correlated with each other, indicating that they may be sensitive to the same hierarchy-related change in neural dynamics. These best-performing features are based on e.g., fluctuation analysis, forecasting, or mean-dependent prediction. However, the computationally simplest among those is the ‘variability of the mean’ (StatAvl100 in Figure 3a, referred to as VarM from hereon), calculated as the standard deviation of the time-series’ mean in non-overlapping moving time windows of 333 ms (Figure 3b). A higher feature value indicates a higher standard deviation of the mean over time, thus a higher variability in the mean across the signal.

**Figure 3:**
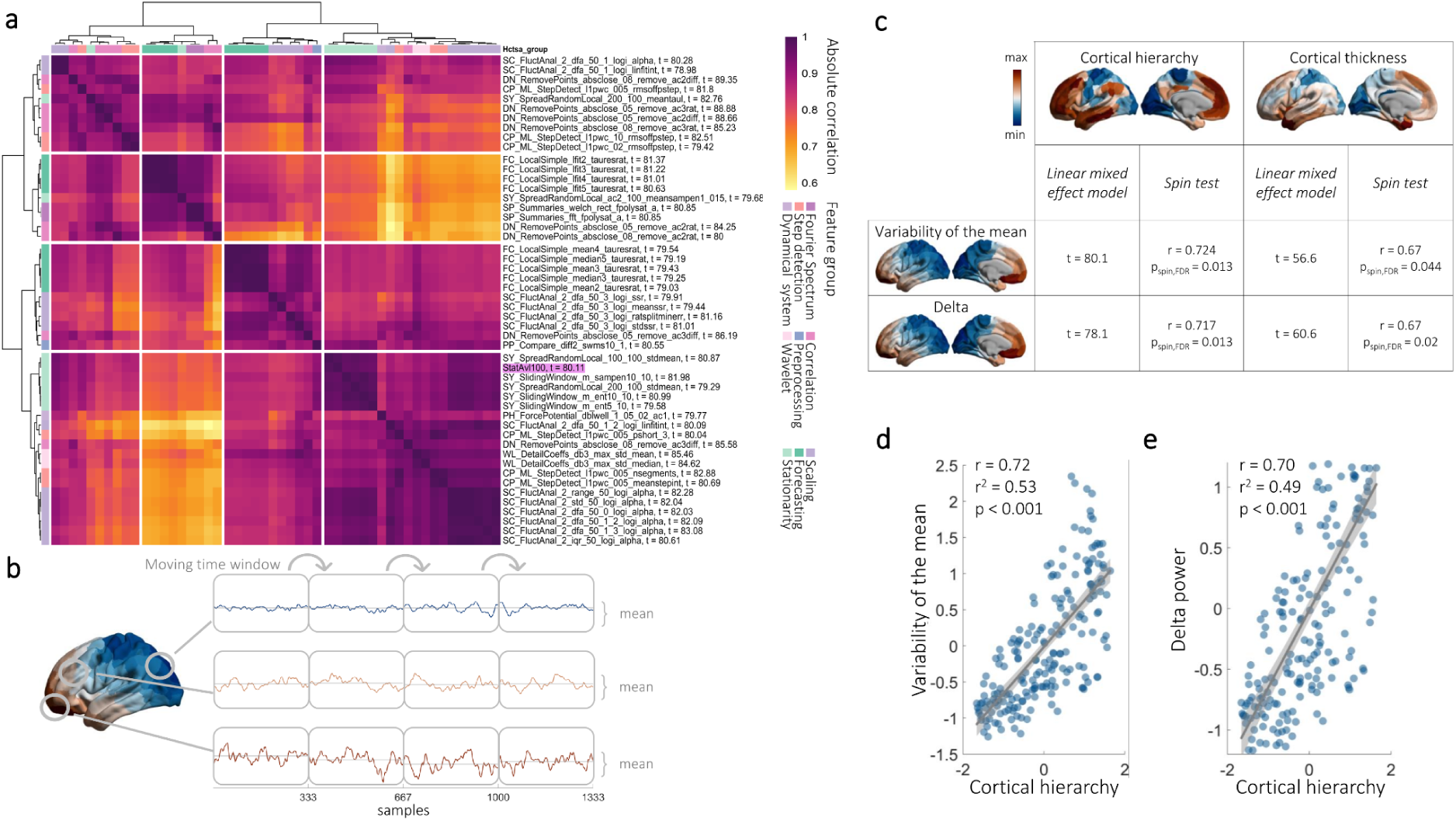
Analysis of neurophysiological time-series features changing along the axis of cortical hierarchy. a: pairwise correlation and cluster analysis including the 50 features showing the most significant change along the axis of the archetypal cortical hierarchy map in feature-wise LMEMs. Many features are highly correlated. The StatAvl100 feature is highlighted as the relationship of this feature with cortical hierarchy was further investigated, however for clarity referred to as VarM. b: Computation of the feature VarM: For each parcel of each participant the time-serie’s mean is iteratively calculated using a non-overlapping moving time window approach with epochs of 333 ms. The VarM feature is defined as the standard deviation of the means calculated across these time windows. Topography is the group-average of the z-scored VarM feature. c: Statistical results from the LMEM and spin test comparing the relationship of the VarM with the archetypal cortical hierarchy topography as well as cortical thickness. It reveals how the VarM feature varies robustly along the archetypal cortical hierarchy map as well as cortical thickness. Additionally, the results from the same analyses investigating how delta power varies systematically along the cortical hierarchy are listed for comparison. d: Scatter plot showing the group-averaged z-scored VarM feature for each parcel of the cortical hierarchy. Parcels situated higher in cortical hierarchy show a higher variability in the mean, thus an increased VarM value. The gray line describes the linear fit for the displayed data with the 95% confidence bounds as gray shaded area. e: Group-averaged z-scored delta power across cortical hierarchy. Delta power linearly increases with cortical hierarchy with highest delta power in parcels high in cortical hierarchy. The gray line describes the linear fit for the displayed data with the 95% confidence bounds as gray shaded area.

The variability of the mean is high in prefrontal and orbitofrontal brain areas and decreases from there towards sensorimotor and occipital brain areas (Figure 3c). Both, LMEM and spin test show that the VarM feature and cortical hierarchy are significantly related. Also, when participant-specific cortical thickness data is used, the effects remain (Figure 3c). Thus, the VarM increases systematically along the axis of cortical hierarchy, also displayed in Figure 3d (for the full LMEM results, see Supplementary Table 4,5). This plot additionally shows that the variability of the VarM feature itself increases from brain areas of low cortical hierarchy towards brain areas high in cortical hierarchy.

The statistically significant increase of the VarM feature along cortical hierarchy is partly independent from the timescale used for computation as different moving time window sizes (83 ms - 3.3 s) result in similar significant effects and spatial distribution of the feature (t_LMEM_ >= 63.4, for more details, see Supplementary Figure S3). However, the strongest association between VarM and hierarchy is observed for a window size of 333 ms. Again, in line with the analysis of delta power, we used surrogate data that preserve spatial autocorrelation and confirmed that results remained significant (Supplementary Figure S1b).

Next to features related to the time-series’ mean, clustering revealed another unrelated feature group (starting with ‘DN_RemovePoints’, Figure 3a). These features compare the autocorrelation between the raw time-series and when at least 50% of sample points are either removed or saturated. Even though, interestingly, features in this group change consistently across the cortical hierarchy it is unclear what aspects of neural dynamics they capture. Thus, further explanation of this feature and feature plots will be only included in the supplements (Supplementary Figure S4).

### Alpha power distribution along the cortical hierarchy changes with age

The presented data set includes participants across a wide age range, thus allowing us to address the relevant question if feature gradients change with age. Therefore, we studied the interaction of age and cortical hierarchy on time-series features in the LMEM.

As in the previous section we start with the traditional features. Similar to the main effect of hierarchy interaction of the archetypal cortical hierarchy axis and age are strongest in low frequency bands (delta: t_LMEM_ = -17.2, theta: t_LMEM_ = -12.1, alpha: t_LMEM_ = 21.4, beta: t_LMEM_ = -5.0, gamma: t_LMEM_ = -11.1). This is still the case when using individual cortical thickness (delta: t_LMEM_ = -14.2, theta: t_LMEM_ = -11.1, alpha: t_LMEM_ = 20.1, beta: t_LMEM_ = -8.1, gamma: t_LMEM_ = -2.7). In particular, for the alpha band, power changed most robustly across age and cortical hierarchy compared to other frequency bands (for full LMEM results, see Supplementary Table S6,7). From a very posterior-anterior power distribution, with highest alpha power in occipital brain areas decreasing towards anterior brain areas, this distribution shifts with age. In older age, strong alpha power spreads additionally to the temporal lobe and decreases towards anterior brain areas (Figure 4a). As in the previous section, we employed a permutation approach for the LMEM to account for confounding effect such as spatial autocorrelations. Removing the relationship between a participants age and the feature distribution along the cortex and retesting results using the LMEMs 10000 times confirmed strong significance for alpha power (for more details, see Supplementary Figure S7).

**Figure 4:**
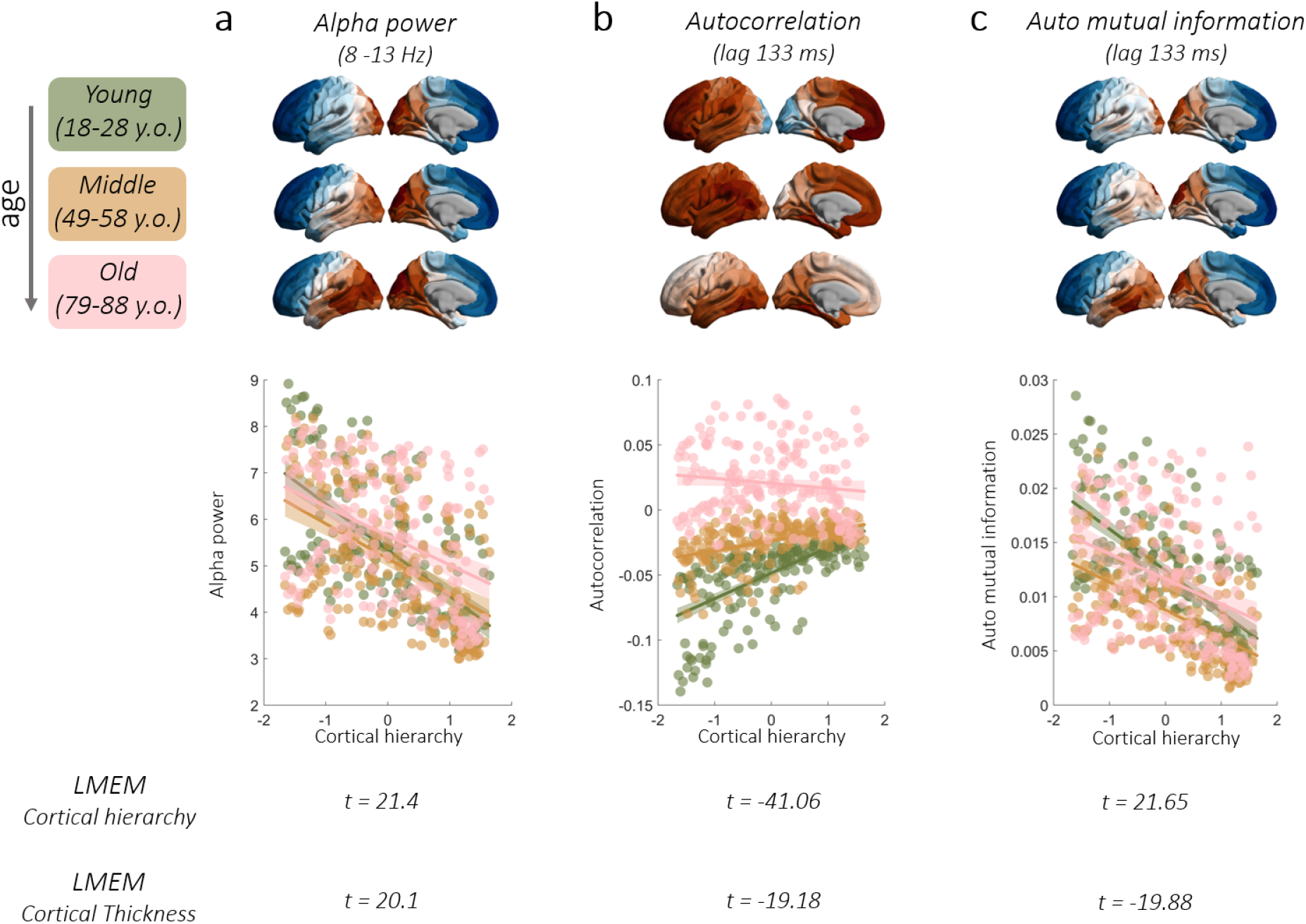
Changes of feature gradients with age. a: alpha power (8-13 Hz), as well as b: autocorrelation and c: auto mutual information (both at lag 133 ms) across age. a: alpha power plotted across the cortical mantle averaged across 50 participants of each age group. Young: 18-28 y.o., middle: 49-58 y.o, old: 79-88 y.o.. While in a young age group the alpha power varies along posterior-anterior gradient with highest alpha power in occipital brain areas, this distribution shifts especially in old age. Here, alpha power in particular increased in occipital and temporal brain areas. Additionally, scatterplots with alpha power plotted against cortical hierarchy, group-averaged across the age groups, show more closely how this change in spatial distribution relates to cortical hierarchy. Young age group in green, middle age group in orange, old age group in pink. The scatter plot shows how the steep decrease of alpha power across cortical hierarchy in a young age group decreases with age leading to a flatter slope in the relationship between alpha power and cortical hierarchy. Finally, the t-values resulting from the LMEMs showing the interaction effect of age and cortical hierarchy or age and cortical thickness, respectively, show the statistical significance of this relationship. b: Inversely to the alpha power gradient, for a young and middle age group, autocorrelation at the lag of 133 ms is lowest in occipital brain areas increasing towards prefrontal areas following a posterior-anterior gradient. Values approaching zero indicate less periodicity in the signal while values diverging from zero indicate increased signal periodicity. Also here, with higher age, this change of periodicity across the cortical hierarchy decreases towards a more uniform distribution of autocorrelation across the cortical hierarchy. c: Auto mutual information (AMI) at a lag of 133 ms, similarly like alpha power, shows a posterior-anterior gradient with increased AMI in the occipital lobe. This distribution shifts with older age and therefore having highest values in occipital brain areas and additionally in the temporal lobe. This shift constitutes a less strong decrease of AMI across cortical hierarchy with age as visible in the scatter plot. For a different representation of the results, see Supplementary Figure S6.

Further, neither the 1/f exponent nor the peak frequency is consistently affected by the interaction of cortical hierarchy and age (1/f: cortical hierarchy and age: t_LMEM_ = 2.7, cortical thickness and age: t_LMEM_ = -9.4; peak frequency: cortical hierarchy and age: t_LMEM_ = 20.7, cortical thickness and age: t_LMEM_ = 1.8).

### The hierarchy-related gradient of autocorrelation and auto mutual information is modulated by age

The LMEM also revealed a spatial change in the cortical distribution of the additionally extracted time-series features related to cortical hierarchy mediated by age. The above-mentioned VarM feature showed significant interactions between hierarchy and age (see Supplementary Table S4). However, other features showed significantly stronger interaction effects. Thus, we pairwise correlated and clustered the 50 features with the highest significant interaction effect of age and the archetypal cortical hierarchy map on the feature value. This results in different subclusters which have high correlations with other features within and less correlation to features outside the cluster (for more details, see Supplementary Figure S5, Table S8). Following our approach from the main analysis of hierarchy, we select autocorrelation and auto mutual information (AMI) as two representative and interpretable features for further analysis.

#### Autocorrelation

One subcluster includes features related to the autocorrelation function of the time-series. Autocorrelation describes how much a shifted version of the time-series correlates with itself and thus provides information about the periodicity and self-similarity of the signal. The autocorrelation features resulting in the highest t-values were found at time lags 83 ms and 87 ms as well as 130 ms and 133 ms. Figure 5a shows the whole autocorrelation function averaged across parcels in different hierarchy levels as well as different age groups (18-28, 49-58, 79-88 years old). It demonstrates how the autocorrelation function differs not only across parcels at different hierarchy levels but, importantly, that age mediates this relationship. This is also depicted by the t-values resulting from the LMEM in the lowest panel of Figure 5a where cortical hierarchy has a very high main effect on autocorrelation for time lags in which also the interaction of age and cortical hierarchy shows significant effects. The effect of the interaction of age and cortical hierarchy is greatest in absolute numbers at a lag of 133 ms (t_LMEM_ = -41.06, for full LMEM results, see Supplementary Table S9). The permutation approach for the LMEM confirmed strong significance for this effect (for more details, see Supplementary Figure S7).

**Figure 5:**
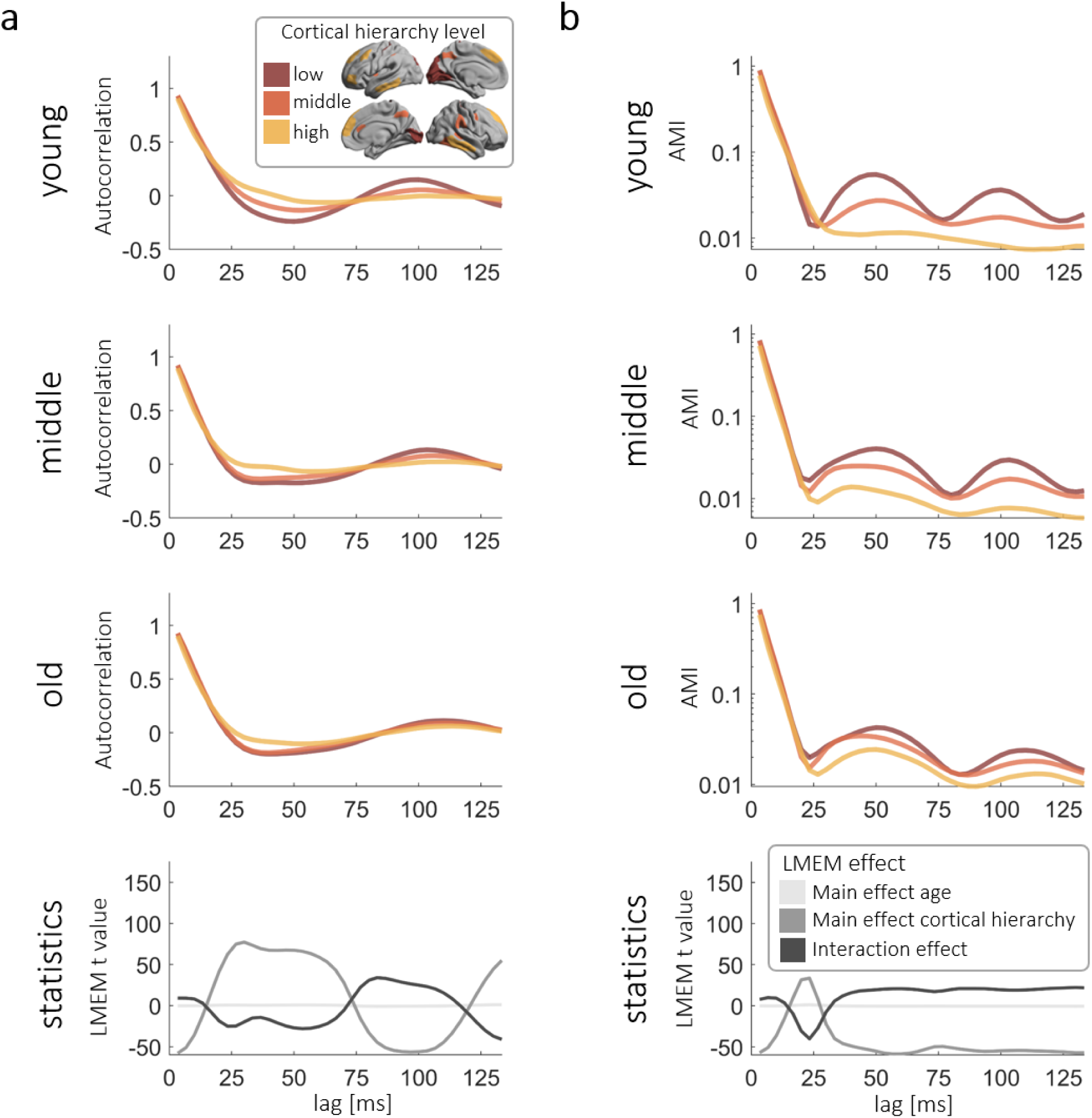
Autocorrelation and auto mutual information (AMI) changes along cortical hierarchy differently across lifespan. a: autocorrelation function for 50 participants of different age groups (18-28, 49-58, 79-88 years old, shown in the panels) and across parcels at different levels of cortical hierarchy (averaged across ten parcels which are lowest, middle, or highest in cortical hierarchy; inserted figure) across different lags. The lowest panel shows the respective t-statistics resulting from the LMEM for the main effect of age, the main effect of cortical hierarchy, and the interaction of both in different gray shades. The latter being strongest in absolute numbers at a lag of 133 ms. b: AMI function plotted across different age groups and parcels at different hierarchy levels (as in a) in a log scale. It shows how the AMI varies across the axis of cortical hierarchy, which changes across age. In the lowest panel, the t-values resulting from the LMEM are shown as described in a.

Taking a more specific look at the spatial distribution of the autocorrelation at the lag of 133 ms (Figure 4b) across the cortex, autocorrelation increases mainly along cortical hierarchy in a young and middle-aged group. In occipital visual brain regions, lower in cortical hierarchy, autocorrelation at this lag is negative and increases towards zero in more frontal brain areas. This is equivalent to a higher periodicity in posterior brain areas at 133 ms, respectively 7.5 Hz, which decreases along cortical hierarchy. In older age, this spatial distribution of autocorrelation diverges from the axis of cortical hierarchy closer to a superior-inferior gradient. Here, autocorrelation is highest in the temporal lobe and occipital visual areas and decreases towards superior frontal brain areas. The statistical effect that was found for the interaction of the archetypal cortical hierarchy and age on autocorrelation at the lag of 133 ms can be replicated when including participant-specific cortical thickness data in the LMEM, however less strongly (t_LMEM_ = -19.18, for full LMEM results, see Supplementary Table S10).

#### Auto mutual information

Another subcluster includes predominantly features related to auto mutual information (AMI) with different computations, different parameters, and lags (for more details, see Supplementary Figure S3). As they all point toward a common underlying neural dynamic, we focus the analysis on the AMI function with different time lags.

AMI describes the statistical dependence between two signals. Compared to the autocorrelation function it captures not only a linear relationship but also non-linear properties. Generally, AMI changes along cortical hierarchy and is additionally mediated by age which is displayed by the t-statistics plotted in the lowest panel of Figure 5b. Most significant is this latter interaction effect at the lag of 23 ms (t_LMEM_ = -40.28). However, to connect our results of the autocorrelation function and the AMI function, we decided to investigate how the spatial distribution of AMI at the lag of 133 ms changes across age. Also here, AMI is affected by the position of a parcel within the axis of cortical hierarchy as well as age (t_LMEM_ = 21.65, for full LMEM results, see Supplementary Table S11).

Here, similar to the autocorrelation, AMI at the lag of 133 ms shows a posterior-anterior gradient in a young and a middle-aged group. However, inverted to autocorrelation, AMI is high in occipital areas of the brain and decreases towards the prefrontal brain areas. In an older age group however, the spatial distribution of AMI resembles autocorrelation with a higher feature value in temporal lobe, decreasing towards more superior frontal brain areas (Figure 4c). The permutation approach for the LMEM confirmed strong significance for this effect (for more details, see Supplementary Figure S7). Using individual cortical thickness-related data instead of the archetypal cortical hierarchy map, this interaction effect can be replicated at the lag of 23 ms (t_LMEM_ = -30.20) as well as 133 ms (t_LMEM_ = 19.88, for full LMEM results, see Supplementary Table S12) however with smaller t-values.

## Discussion

The dynamics of neurophysiological brain activity vary between different brain areas and are constrained by their underlying anatomical properties, some of which change systematically across the cortical hierarchy. In this study, we aimed to identify features of spontaneous brain activity that change consistently across this axis of cortical hierarchy. For this analysis, we did not only leverage traditional time-series features related to the power spectrum of the signal, but we additionally performed massive time-series feature extraction ^55,56^. By including participant-specific spatial information on the distribution of single time-series features along the cortex, we report individual features that reflect cortical hierarchy across individuals. We were able to identify spectral power in the slow frequency bands and the variability of the mean’s signal as neurophysiological markers of first, generalized archetypal cortical hierarchy ^8^, and second, individual cortical thickness serving as a proxy for cortical hierarchy. Additionally, by including participants with a wide age range, we reveal that simple time-series features not only vary along cortical hierarchy but further systematically capture age-related changes in cortical hierarchies.

Traditionally, features related to the power spectrum have been used to characterize resting-state brain activity. This includes spectral power in canonical frequency bands but also extends to the 1/f exponent of the aperiodic component of the power spectrum as well as the peak frequency of alpha. The 1/f exponent and alpha peak frequency have previously been linked to the posterior-anterior gradient as well as cortical thickness ^23^. Importantly, our comprehensive analysis reveals other features with a significantly stronger association with cortical hierarchy. Specifically, spectral power in the delta frequency band describes cortical hierarchy most robustly among all traditional features included here. Previously, endogenous delta band brain activity has been reported during rest ^60^ and shown to be strongest in frontal brain areas ^61,62^. However, our results provide the first evidence that delta power changes gradually across the cortex in a spatially specific manner that significantly follows the cortical hierarchy.

However, spectral features have shortcomings. They might be skewed by non-rhythmic brain activity ^63^ or sudden bursts of activity ^63^. Computation of the power spectrum using the Fourier transform assumes stationarity and a sinusoidal waveform shape ^64,65^ which are often violated by brain activity. Taken together, many potentially important aspects of brain activity are not captured by spectral features. Indeed, our comprehensive time-series phenotyping reveals a number of features that are proxies of cortical hierarchy, some of which even outperform delta power. This confirms recent findings that particular time-series features can outperform spectral features, for example at predicting different brain states ^66^ or age ^67^.

Among the features displaying a gradient along the cortical hierarchy, the feature ‘variability of the mean’ (VarM) stands out, due to its interpretability on the one hand and high correlation with other features relating to the signal’s mean on the other hand. This feature captures the variability of the signals’ mean over time, increasing linearly with cortical hierarchy with the highest variability at the apex of the hierarchy. This increased variability in the signal might reflect the need to integrate diverse inputs leading to more dynamic and adaptive, context-dependent activity patterns at areas situated higher in cortical hierarchy. Conversely, regions lower in hierarchy exhibit lower mean signal variability thereby ensuring possibly more context-independent, stable processing patterns necessary to reliably react to the surrounding environment. Thus, higher variability in regions higher in cortical hierarchy should not be disregarded as noise but is rather a substrate for the intra- and interindividual differences in behavior ^68,69^. This result resonates with other studies, which described how the interindividual variability is also increased in the BOLD signal ^70^ as well as functional connectivity ^71^ in brain areas situated high in cortical hierarchy. This spatial heterogeneity is also described for measures of structure-function coupling which decreases with cortical hierarchy ^12,72,73^, possibly facilitating a higher variability in the functional outcome ^74^ of brain areas high in cortical hierarchy. The phenomenon might be scaffolded by the spatial distribution of intracortical myelin. As myelin is low in areas higher up in cortical hierarchy ^10,13^, it gives room for synapses and synaptic spines to emerge and disappear and thus remodel these brain areas to a higher degree than brain areas low in hierarchy. Here, higher levels of myelin decreases the possibility of synaptic formation ^75^, decreasing variability in these regions.

Naturally, variability of the mean and spectral power are related measures, both describe deflections in the signal. Importantly, the window length for computation of the variability of the mean varied between 25 and 1000 samples, thus spanning 83 ms up to >3 s. All these measures show an increase of the mean’s variability with increasing cortical hierarchy and the same spatial pattern across the cortex. Thus, the variability in the mean cannot be accounted for by power in specific frequency bands. Additionally, as described before, unlike the Fourier transform, the feature does not assume stationarity of the brain signal but rather adapts to an inherent non-stationarity of brain signals and their non-sinusoidal waveform shapes that change across the cortex ^63–65^. Given these observations, the power and mean variability are related but probe different characteristics of the signal. Therefore, this measure of variability can be an important complementary neurophysiological feature to characterize cortical hierarchy.

Crucially, this study includes a large participant cohort with a wide age range. Including individual structural and functional data as well as age of the participants in the analysis, promotes the understanding how time-series features showing a gradient along the cortical hierarchy might additionally change with age. Resulting from the performed analyses, we found three measures showing robust spatial changes of their feature distribution across cortical hierarchy with age, namely alpha power, autocorrelation, and auto mutual information. Even though the lags where we found the strongest effect for autocorrelation and auto mutual information (∼7.5 Hz) are at the lower end of the classical alpha frequency band, all three measures show similar age-related changes.

Age-related changes of alpha power are characterized by a focal distribution of highest alpha power in occipito-parietal areas that progressively spreads anterior along the frontal lobe with increasing age. In relation to cortical hierarchy, this means that the change of alpha power across cortical hierarchy decreases with age. This relates to studies that show alpha power decreases with age ^40,51^ in which the decrease in power is strong in occipital brain areas and decreases when going to more frontal brain areas ^40,50^.

Age-dependent changes of alpha across the cortical hierarchy are similar to age-related changes in autocorrelation. Autocorrelation captures linear self-similarity of the signal as well as periodicity and shows a clear change along the cortical hierarchy with age. This is to be expected, as mathematically the autocorrelation function is simply the Fourier transform of the power spectrum. Also, other studies demonstrated that the autocorrelation function changes across cortical hierarchy. The first zero crossing ^76^ and the timescale resulting from the autocorrelation function ^26,27^ vary along the axis of cortical hierarchy. Gao et al. further showed that the timescale averaged across the cortex shortens with age ^26^. We extend this finding by describing how the whole autocorrelation function varies across cortical hierarchy and age. Particularly at the lags, where cortical hierarchy has an effect on the autocorrelation function, age mediates this relationship. However, the spatial topography does not change for all lags uniformly with age but rather differently. Specifically for autocorrelation at a lag of 133 ms or 7.5 Hz, highest periodicity is observed in the occipital brain areas for young and middle aged groups which then diminishes with age until cortical hierarchy hardly affects the distribution of autocorrelation across the cortex in a group of old age.

Importantly, next to the autocorrelation also auto mutual information shows age-related changes along cortical hierarchy. This measure captures, in contrast to autocorrelation, additionally nonlinear dependencies ^77^ and is not inherently related to the power spectrum. Particularly, auto mutual information at a lag of 133 ms (7.5 Hz) shows the highest values in occipital brain areas, which with age shifts towards including temporal brain areas as well. Hence also here, we find that the change of a measure of self-similarity of the signal across cortical hierarchy decreases with age, thus describing similar aging-specific changes as alpha power and autocorrelation at lag 133 ms. Following this line of thought, the auto mutual information possibly includes complementary information to what can be captured only by the power spectrum and autocorrelation. Additionally, it gives the idea that the linear and nonlinear dependencies capture similar microstructural changes across the axis of cortical hierarchy, which are in turn mediated by age possibly in a similar manner.

Even though this study benefits from participant specific structural and functional data, age variability, and a large sample size, it does not come without limitations. We specifically tailored the analysis to the identification of gradients across the cortex, thereby becoming insensitive to age-related regional changes that have been analyzed and reported elsewhere ^40,78^. Further, although we included two different measures of cortical hierarchy, additional features of this axis have been reported, for instance the first functional gradient ^9^ as well as its intersubject variability ^71^, T1-weighted/T2-weighted MRI ratio ^79^, or evolutionary expansion ^80^, which are openly accessible ^57^. However, we decided to implement the archetypal map as a measure of cortical hierarchy, as it already captures information of many structural, functional, developmental, and evolution-related hierarchy measures ^8^. Further we decided on a complementary approach in which we performed analyses using individual cortical thickness data, to show the alignment of a participant-specific anatomical hierarchy-sensitive measure and the individual functional time-series.

In conclusion, this study shows how slow spectral power as well as other simple time-series measures, such as the variability of the mean, reflect cortical hierarchy and thus could serve as a neurophysiological marker of health and disease. Altered functional gradient of hierarchical organization using fMRI has been associated with e.g., schizophrenia ^53,81^ and autism ^82^. Biological and structural changes in these disorders have also been linked with the first principal component of gene expression, which in turn follows cortical hierarchy ^83^. Following this line of thought, also neurophysiological measures of cortical hierarchy, such as the variability of the mean, could possibly be extracted from simple EEG measurements and serve as a marker of neurological disorders. Further, this study highlights that characteristics of neural dynamics vary along the cortical hierarchy differently across the lifespan. In literature, the change of other brain activity measures along the hierarchical gradient across development, adolescence and aging is already described. For instance, the gradient capturing the highest variability in functional connectivity, spanning from the unimodal to transmodal axis, emerges only during adolescence ^52,53^ and disperses during aging ^54^. However, future work will have to clarify the exact determinants of changes of brain activity along the cortical hierarchy and across the lifespan. Our comprehensive time-series phenotyping is a crucial step in this direction.

## Materials and Methods

### Dataset

The data used in this study was published by the Cambridge Centre for Ageing and Neuroscience (CamCAN data repository) and contains data from three study stages with various measures such as home interviews, structural as well as functional brain imaging, and cognitive assessments ^84,85^. The following data selection and (pre)processing steps follow analyses as published in ^40,85^.

Originally, the repository encompasses data of 650 participants, of which the structural T1 and T2 MRIs as well as MEG resting-state data (eyes-closed) of 450 participants were used for further analysis. Due to motion artifacts, low signal-to-noise ratio, sleep during the resting state, or unsuccessful Freesurfer reconstruction, 72 participants were excluded. Out of the remaining datasets, 350 were randomly selected following a balanced design: within an age range from 18 to 88, 50 participants were selected for each 10-year age range, balanced in sex. Participants were excluded if they were cognitively impaired, had communication difficulties, or had medical problems based either on self-reports or diagnosis (for more details, see ^85^).

The study was conducted following the Declaration of Helsinki (World Medical Association, 2013) and approved by the local ethics committee, Cambridgeshire 2 Research Ethics Committee.

### MRI data acquisition

Anatomical imaging was performed with a 3T Siemens TIM Trio scanner equipped with a 32-channel head coil. T1-weighted images were obtained from 3D MPRAGE sequences with TR = 2250 ms, TE = 2.99 ms, TI = 900 ms, FA = 9 degrees, FOV = 256 x 240 x 192 mm, 1 mm isotropic resolution, GRAPPA = 2, TA = 4.5 min. T2-weighted images were acquired using 3D SPACE sequences with a TR = 2800 ms, TE = 408 ms, TI = 900 ms, FOV = 256 x 256 x 192 mm, 1 mm isotropic resolution, GRAPPA = 2, TA = 4.5 minutes.

### MEG data acquisition

At the MRC Cognition and Brain Science Unit, University of Cambridge, UK, MEG recordings were conducted in a magnetically shielded room. The facility is equipped with a 306-channel VectorView MEG system manufactured by Elekta Neuromag in Helsinki. This system comprises 102 magnetometers and 204 planar gradiometers. The data was sampled at 1 kHz and a high pass filtered at 0.03 Hz. During MEG recordings, participants were seated for 8 minutes and 40 seconds with instructions to keep their eyes closed and remain still. Head position was monitored using 4 coils, while electrocardiography (ECG) and electrooculography (EOG) were employed to measure heart activity as well as horizontal and vertical eye movements.

### MRI processing

In order to reconstruct cortical brain activity based on MEG sensor level data, both T1 and T2-weighted images were processed using Freesurfer software 6.0.0 (https://surfer.nmr.mgh.harvard.edu/). Surface-based mapping ^86^ was applied, which resampled individual cortical surfaces to 1,002 vertices per hemisphere with a scaling factor of 10, based on the ’fsaverage’ template mesh (afni.nimh.nih.gov/download/) supplied by Freesurfer and SUMA. Individual meshes were aligned to the Neuromag sensor space using anatomical landmark coordinates provided by Cam-CAN. The "single shell" method, implemented in Fieldtrip, was utilized to compute the lead fields and individual head models for MEG source projection.

### MEG processing

The CamCAN repository offers preprocessed data, which included removal of external noise and artifacts, line-noise suppression (50 Hz and its harmonics), and corrections for noisy channels and head movements. For this purpose, Elekta Neuromag Maxfilter 2.2 was applied using temporal signal space separation (10-second sliding window, 0.98 correlation limit) to the continuous MEG data ^84^.

The data was further preprocessed using Fieldtrip ^87^ including downsampling to 300 Hz, high pass filtering at 1 Hz (1st order Butterworth filter), and epoching to 10 s trials. Then, automatic artifact detection was used to remove trials containing muscle artifacts, following https://www.fieldtriptoolbox.org/tutorial/automatic_artifact_rejection/. In detail, the data was bandpass filtered between 110 - 140 Hz using a 9th-order Butterworth filter. For each channel, the signal amplitude was calculated (Hilbert envelope) and subsequently z-transformed. These z-values were averaged across channels resulting in global averages for every time-point. Time points exceeding a z-threshold of 14 were marked as part of an artifact, and the time-period of the artifact additionally padded to both sides. Trials containing artifacts were rejected.

Then, the data were low-pass filtered at 70 Hz using a 1st-order Butterworth filter and independent component analysis (ICA) was applied in order to correct for ocular and cardiac artifacts. Ocular artifacts were identified if the average coherence with the EOG was > 0.3 and the amplitude correlation coefficient between the signals was > 0.4. Cardiac artifacts were corrected based on the average coherence with the ECG (coherence > 0.3) or on the averaged maximum peaks time-locked to the ECG (QRS complex,https://www.fieldtriptoolbox.org/example/use_independent_component_ analysis_ica_to_remove_ecg_artifacts/). If the signal-to-noise ratio was very low in the EOG/ECG, respective ICA components had to be manually selected. For each participant, all selected ICA components as well as the resulting, cleaned data were additionally visually checked for quality control.

To ensure participants were awake, the vigilance was rated according to the American Academy of Sleep Medicine (https://aasm.org/), and 30 ‘awake’ trials per participant were randomly selected (> 50% alpha activity within a trial). Finally, individual data derived from the magnetometers (102 channels) was source-reconstructed, by projecting the sensor level signals onto the source space of the cortex using unit-noise gain linear constrained minimum variance (LCMV) beamforming ^88^. Spatial filters were created (with a regularization lambda of 5% and a free-orientation forward solution) and applied to the sensor time-series data. This process yielded source-level time-series data for 200 regions defined according to the Schäfer atlas ^89^. MEG preprocessing was performed in Matlab (R2018b).

### Cortical thickness estimation

For each cortical vertex, cortical thickness was estimated using T1-weighted MRI data, measuring the distance from the gray-white matter boundary to the pial surface. Smoothing was performed using a heat kernel with a full width at half maximum of 12 mm in AFNI (https://afni.nimh.nih.gov/). One participant was excluded from all analyses including cortical thickness related data as an outlier. The correlation of this participant’s cortical thickness data had a difference of more than ten standard deviations from the mean cortical thickness and additionally more than five standard deviations difference to the mean cortical thickness of the respective age group.

### Time-series feature computation

In order to deeply phenotype the time-series, features were extracted using the highly comparative time-series analysis toolbox (hctsa, ^55,56^ using Matlab R2018b). The computation was performed on 30 trials of 10 s length, and averaged across trials, resulting in 7525 features for 200 parcels and 350 participants. Features with constant or complex values, outliers, or invalid numbers were removed leading to 5961 remaining features for further analysis, encompassing measures related to autocorrelation, fluctuation analyses, distribution, time-series model fitting, forecasting, and many more. Additionally, more commonly used features were calculated, such as spectral power in canonical frequency bands, the exponent of the 1/f decay of the aperiodic component, and individual alpha peak frequency. Spectral power was computed through frequency analysis employing multitapers based on discrete prolate spheroidal sequences (DPSS) with 1 Hz smoothing using Fieldtrip ^87^ using Matlab (R2022b). Then, power was averaged across canonical frequency bands (delta: 1-3 Hz, theta: 4-7 Hz, alpha: 8-13 Hz, beta: 14-30 Hz, gamma: 31 - 70 Hz) and normalized by dividing by the sum of the whole power spectrum. To compute the aperiodic (1/f exponent or slope) component of the MEG signals, we passed the source power (1-60 Hz) to the specparam algorithm ^59^ using the following settings: peak width limits [1-8]; maximum number of peaks: infinitive; minimum peak height: 0; and aperiodic mode: fixed. Moreover, we estimated alpha peak frequency using the center of gravity method ^90^, which estimates the weighted average of frequencies in the alpha band range weighted by their power. Before further analyses, all features were z-scored across parcels for each participant.

### Global cortical hierarchy maps

Using the neuromaps toolbox ^57^, the archetypal cortical hierarchy map according to Sydnor et al., ^8^ was obtained. This sensorimotor-to-association axis is revealed by macroscale cortical gradients across different measures and species. It is congregated from ten different measures: T1-weighted to T2-weighted ratio ^91^, the first principal gradient of functional connectivity ^9^, evolutionary cortical expansion ^80^, allometric scaling ^92^, aerobic glycolysis ^93^, cerebral blood flow ^94^, first principal component of cortical gene expression ^11^, NeuroSynth ^95^, externopyramidization ^96^, and cortical thickness (Human Connectome Project S1200 data). For each data type, values for each left hemispheric parcel ^79^ were ranked. Parcel-wise averaging across all ten measures results in one archetypal generalized measure of cortical hierarchy.

### Linear mixed-effect modeling

As this analysis aims to characterize individual MEG features and their distribution across the cortex, no dimensionality reduction techniques are employed. Instead, each time-series feature was analyzed individually. As the brain shows interindividual differences and further changes throughout life in structural and functional measures, we included the individual participant and age as factors in the analysis. To this end, we chose linear mixed effect models (LMEMs) as they are able to incorporate fixed effects of cortical hierarchy and age as well as their interaction and random effects of the individual participant:

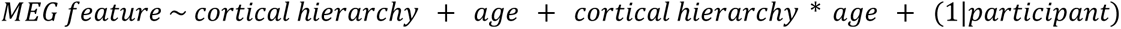

As the archetypal cortical hierarchy is an over-subject generalized concept ^8^, this analysis may miss deviations due to variability in the structure and function of the individual. Thus, individual cortical thickness measures were extracted from the structural imaging included in the CamCAN repository which can serve as a participant-specific proxy for cortical hierarchy ^13^ and which is known to change during lifespan ^31,32,40^. Thus, we use the same LMEM approach, with the features we found to have highly significant effects, and retest them using individual cortical thickness instead of archetypal cortical hierarchy. The analyses described here were performed using Matlab (R2022b).

### Clustering

Many of the hctsa features are highly correlated among each other, especially when they pick up on the same neuronal dynamic, estimated using slightly different parameters. Therefore, 50 features with the highest effects in the LMEM were selected and pair-wise correlation analysis was conducted on a participant-specific level using Pearson correlation coefficients. Then, cluster analysis was performed using the Ward.D2 method, which aims to identify compact, spherical clusters. This method implements Ward’s minimum variance criterion ^97^, which involves squaring the dissimilarities before updating the clusters (https://www.rdocumentation.org/packages/stats/versions/3.6.2/topics/hclust). These analyses were carried out using RStudio 2023.03.0. Following cluster analysis, representative features of the subclusters were selected. This selection process aimed to identify features that were both representative of the cluster and relatively simple in their computation and interpretation as the hctsa toolbox encompasses a wide array of features, some of which are non-intuitive or very complex in their computation but highly related to simpler measures. Further analysis was then carried out using these selected features.

### Creating Null models

The following analyses were performed using Python3.11. To provide further complementary analysis to the LMEMs, spin tests were used to quantitatively compare the group-averaged features of interest to cortical hierarchy and cortical thickness maps, respectively. To this end, the neuromaps toolbox ^57^ was employed to obtain an archetypal cortical hierarchy map ^8^, which is subsequently parcellated to the Schäfer200 atlas ^89^ in fsLR32k space, as described before.

To assess the significance of a correlation between time-series features, showing a significant effect in previous analyses, null models were generated, preserving the spatial autocorrelation of the data ^58^, 50.000 permutations) and spin tests were computed. The large number of permutations was chosen to ensure a robust estimation of statistical significance, as the number of spins dictates the minimum possible p-value. Resulting p-values were FDR-corrected to account for multiple comparisons. Since the hierarchical organization of cortical areas develops mostly along the posterior-anterior axis the spin test will be rather conservative since surrogates that are rotations along the posterior-axis still largely preserve the original hierarchy and cannot be considered as pure surrogates.

## Data and code availability

Raw data were provided by the Cam-CAN project and are available at https://camcan-archive.mrc-cbu.cam.ac.uk/dataaccess/ under specified conditions. Single-subject feature values can be downloaded from https://osf.io/h43mz/. All code used for the preprocessing, statistical analyses, and visualization of the results have been deposited in https://github.com/janafhrng/HierarchyCorrelates. Additional z-scored feature matrices are deposited at osf.io/hf7cm. All analyses were conducted using MATLAB R2018b, R2022b, Python3.11, RStudio 2023.03.0. Visualization of features values along the human cortex were performed using the brainspace toolbox (0.1.4.) in Matlab R2022b ^98^ and the scientific color maps 8.0.1 ^99^.

## Acknowledgements

Data collection and sharing for this project was provided by the Cambridge Center for Ageing and Neuroscience (CamCAN). Cam-CAN funding was provided by the UK Biotechnology and Biological Sciences Research Council (grant number BB/H008217/1), together with support from the UK Medical Research Council and the University of Cambridge, UK. This work was further supported by the German Research Foundation (DFG; GR 2024/11-1 to JG; FO 750/5–1 to N.K.F.). We acknowledge support by the Open Access Publication Funds of the University of Münster. We are grateful to Professor Ben Fulcher and Nicolas Chalas for valuable discussions on the methods and results.

## Author contributions

Conceptualization J.G., J.F.

Methodology J.G., C.S., J.F., E.B., N.F.

Investigation J.F., C.S., J.G.

Writing – original draft J.F.

Writing – review & editing J.G., C.S., E.B., N.F., U.D.

Visualization J.F.

Funding acquisition N.F., J.G.

## Corresponding author

Correspondence to Jana Fehring.

