## Supplementary material for "Neurophysiological correlates of cortical hierarchy across the lifespan"

Supplementary material
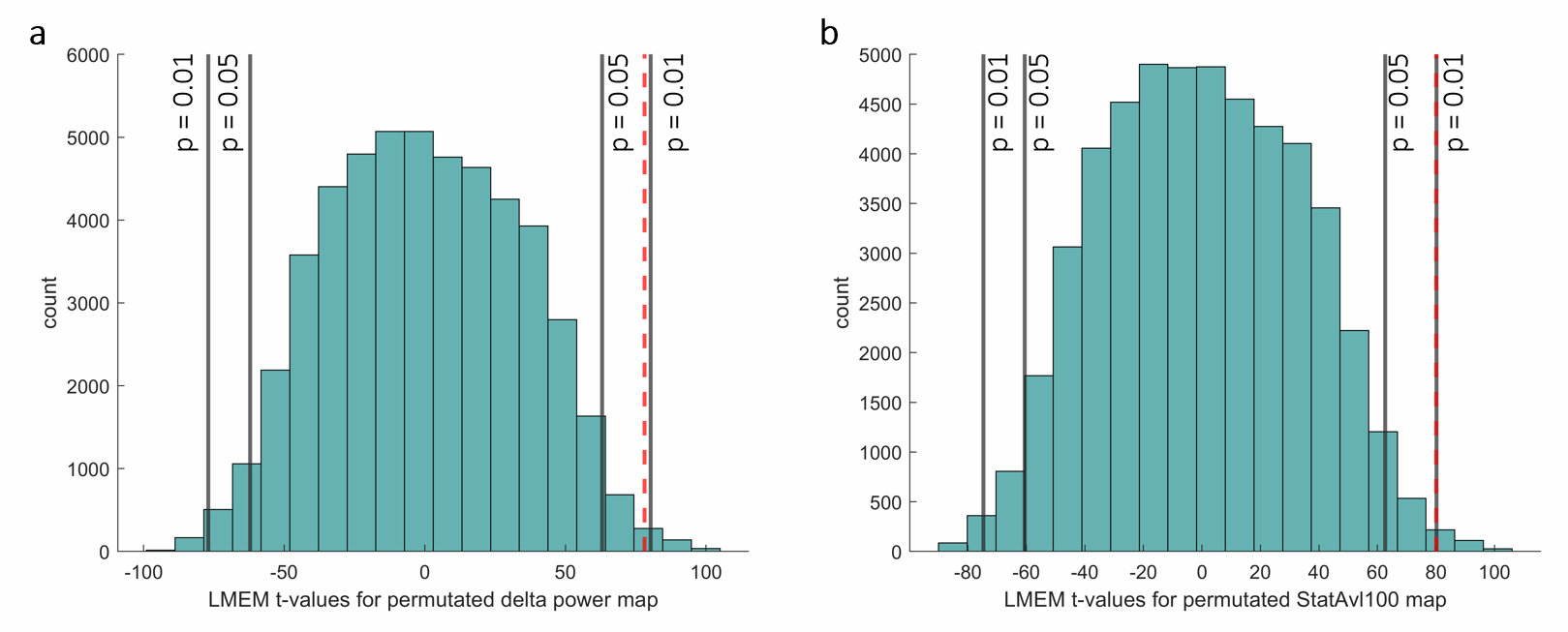

*Figure S1: Distribution of expectable t-values of the permuted time-series feature map (a: delta power, b: StatAvl100 or VarM) in LMEM for a main effect of cortical hierarchy. 50000 permutations were created according to the approach by Alexander Bloch, retaining spatial autocorrelation of the data. With all permuted feature maps LMEMs were recomputed and t-values for the main effect of cortical hierarchy extracted. These t-values create a normal distribution giving the possibility to assign a statistical p-value for the t-value of the LMEM. Gray lines mark the assigned p-values. The red dotted line marks the respective t-value for a: the delta power and b: stationarity of the mean.*

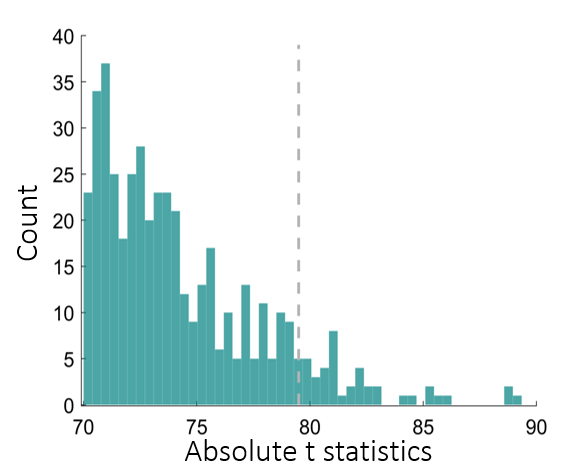

*Figure S2: Distribution of absolute t-values in the LMEM for the main effect of cortical hierarchy for all hctsa features with absolute t-values > 70. Dashed line is the threshold for the 50 top features which were included in further correlation and cluster analysis.*

| **Feature name** | **t-value LMEM** |
| --- | --- |
| DN_RemovePoints_absclose_08_remove_ac2diff | -89.346 |
| DN_RemovePoints_absclose_05_remove_ac3rat | 88.881 |
| DN_RemovePoints_absclose_05_remove_ac2diff | -88.655 |
| DN_RemovePoints_absclose_05_remove_ac3diff | -86.188 |
| DN_RemovePoints_absclose_08_remove_ac3diff | -85.581 |
| WL_DetailCoeffs_db3_max_std_mean | -85.457 |
| DN_RemovePoints_absclose_08_remove_ac3rat | 85.231 |
| WL_DetailCoeffs_db3_max_std_median | -84.621 |
| DN_RemovePoints_absclose_05_remove_ac2rat | 84.246 |
| SC_FluctAnal_2_dfa_50_1_3_logi_alpha | 83.076 |
| CP_ML_StepDetect_l1pwc_005_nsegments | -82.875 |
| SY_SpreadRandomLocal_200_100_meantaul | 82.759 |
| CP_ML_StepDetect_l1pwc_10_rmsoffpstep | 82.512 |
| SC_FluctAnal_2_range_50_logi_alpha | 82.277 |
| SC_FluctAnal_2_dfa_50_1_2_logi_alpha | 82.094 |
| SC_FluctAnal_2_std_50_logi_alpha | 82.043 |
| SC_FluctAnal_2_dfa_50_0_logi_alpha | 82.033 |
| SY_SlidingWindow_m_sampen10_10 | 81.98 |
| CP_ML_StepDetect_l1pwc_005_rmsoffpstep | 81.803 |
| FC_LocalSimple_lfit2_tauresrat | -81.374 |
| FC_LocalSimple_lfit3_tauresrat | -81.217 |
| SC_FluctAnal_2_dfa_50_3_logi_ratsplitminerr | 81.158 |
| FC_LocalSimple_lfit4_tauresrat | -81.012 |
| SC_FluctAnal_2_dfa_50_3_logi_stdssr | -81.012 |
| SY_SlidingWindow_m_ent10_10 | 80.988 |
| SY_SpreadRandomLocal_100_100_stdmean | 80.869 |
| SP_Summaries_welch_rect_fpolysat_a | -80.846 |
| SP_Summaries_fft_fpolysat_a | -80.846 |
| CP_ML_StepDetect_l1pwc_005_meanstepint | 80.686 |
| FC_LocalSimple_lfit5_tauresrat | -80.629 |
| SC_FluctAnal_2_iqr_50_logi_alpha | 80.609 |
| PP_Compare_diff2_swms10_1 | -80.548 |
| SC_FluctAnal_2_dfa_50_1_logi_alpha | 80.28 |
| StatAvl100 | 80.109 |
| SC_FluctAnal_2_dfa_50_1_2_logi_linfitint | -80.086 |
| CP_ML_StepDetect_l1pwc_005_pshort_3 | -80.04 |
| DN_RemovePoints_absclose_08_remove_ac2rat | 79.996 |
| SC_FluctAnal_2_dfa_50_3_logi_ssr | -79.911 |
| PH_ForcePotential_dblwell_1_05_02_ac1 | 79.766 |
| SY_SpreadRandomLocal_ac2_100_meansampen1_015 | 79.677 |
| SY_SlidingWindow_m_ent5_10 | 79.583 |
| FC_LocalSimple_mean4_tauresrat | -79.537 |
| SC_FluctAnal_2_dfa_50_3_logi_meanssr | -79.44 |
| FC_LocalSimple_mean3_tauresrat | -79.427 |
| CP_ML_StepDetect_l1pwc_02_rmsoffpstep | 79.424 |
| SY_SpreadRandomLocal_200_100_stdmean | 79.286 |
| FC_LocalSimple_median3_tauresrat | -79.247 |
| FC_LocalSimple_median5_tauresrat | -79.191 |
| FC_LocalSimple_mean2_tauresrat | -79.031 |
| SC_FluctAnal_2_dfa_50_1_logi_linfitint | -78.985 |
| SC_FluctAnal_2_dfa_50_1_logi_linfitint | -78.985 |

*Table S1: List of the 50 hctsa time-series features with the highest absolute t-values in a LMEM for a main effect of the generalized archetypal cortical hierarchy map. Names and t-values are listed, ranked by absolute t-value.*

| ***Name*** | ***Estimate*** | ***SE*** | ***tStat*** | ***DF*** | ***pValue*** | ***Lower*** | ***Upper*** |
| --- | --- | --- | --- | --- | --- | --- | --- |
| ***(Intercept)*** | *-0.01890632* | *0.00885589* | *-2.13488635* | *69996* | *0.03277376* | *-0.03626385* | *-0.00154879* |
| ***age*** | *5.1827E-05* | *0.00015685* | *0.33042882* | *69996* | *0.74107696* | *-0.00025559* | *0.00035925* |
| ***hierarchy*** | *0.74130173* | *0.00948828* | *78.1281127* | *69996* | *0* | *0.72270471* | *0.75989874* |
| ***age:hierarchy*** | *-0.00203208* | *0.00016805* | *-12.092344* | *69996* | *0* | *-0.00236146* | *-0.00170271* |

*TableS2: Full statistical results for the LMEM exploring the effects of age, cortical hierarchy, the interaction of both as well as individual random effects on delta power.*

| ***Name*** | ***Estimate*** | ***SE*** | ***tStat*** | ***DF*** | ***pValue*** | ***Lower*** | ***Upper*** |
| --- | --- | --- | --- | --- | --- | --- | --- |
| ***(Intercept)*** | *-4.29461241* | *0.06752438* | *-63.6009123* | *69098* | *0* | *-4.42696008* | *-4.16226473* |
| ***age*** | *0.01922962* | *0.0011511* | *16.7054005* | *69098* | *0* | *0.01697346* | *0.02148578* |
| ***thick*** | *1.46695401* | *0.02429613* | *60.3781053* | *69098* | *0* | *1.41933365* | *1.51457438* |
| ***age:thick*** | *-0.00472074* | *0.00042406* | *-11.1321436* | *69098* | *0* | *-0.0055519* | *-0.00388957* |

*TableS3: Full statistical results for the LMEM exploring the effects of age, cortical thickness, the interaction of both as well as individual random effects on delta power.*

| ***Name*** | ***Estimate*** | ***SE*** | ***tStat*** | ***DF*** | ***pValue*** | ***Lower*** | ***Upper*** |
| --- | --- | --- | --- | --- | --- | --- | --- |
| ***(Intercept)*** | *-0.0192026* | *0.0087722* | *-2.1890250* | *69996* | *0.0285983* | *-0.0363962* | *-0.0020091* |
| ***age*** | *0.0000522* | *0.0001554* | *0.3358846* | *69996* | *0.7369589* | *-0.0002523* | *0.0003567* |
| ***hierarchy*** | *0.7529203* | *0.0093987* | *80.1093663* | *69996* | *0.0000000* | *0.7344990* | *0.7713417* |
| ***age:hierarchy*** | *-0.0020461* | *0.0001665* | *-12.2920043* | *69996* | *0.0000000* | *-0.0023724* | *-0.0017199* |

*TableS4: Full statistical results for the LMEM exploring the effects of age, cortical hierarchy, the interaction of both as well as individual random effects on the stationarity of the mean.*

| ***Name*** | ***Estimate*** | ***SE*** | ***tStat*** | ***DF*** | ***pValue*** | ***Lower*** | ***Upper*** |
| --- | --- | --- | --- | --- | --- | --- | --- |
| ***(Intercept)*** | *-4.03609684* | *0.06732278* | *-59.9514268* | *69098* | *0* | *-4.16804938* | *-3.9041443* |
| ***thick*** | *1.3709068* | *0.02422359* | *56.5938821* | *69098* | *0* | *1.32342861* | *1.41838499* |
| ***age*** | *0.013709* | *0.00114767* | *11.9451219* | *69098* | *0* | *0.01145958* | *0.01595842* |
| ***thick:age*** | *-0.00263678* | *0.0004228* | *-6.23649995* | *69098* | *4.5005E-10* | *-0.00346546* | *-0.00180809* |

*TableS5: Full statistical results for the LMEM exploring the effects of age, cortical thickness, the interaction of both as well as individual random effects on the stationarity of the mean.*

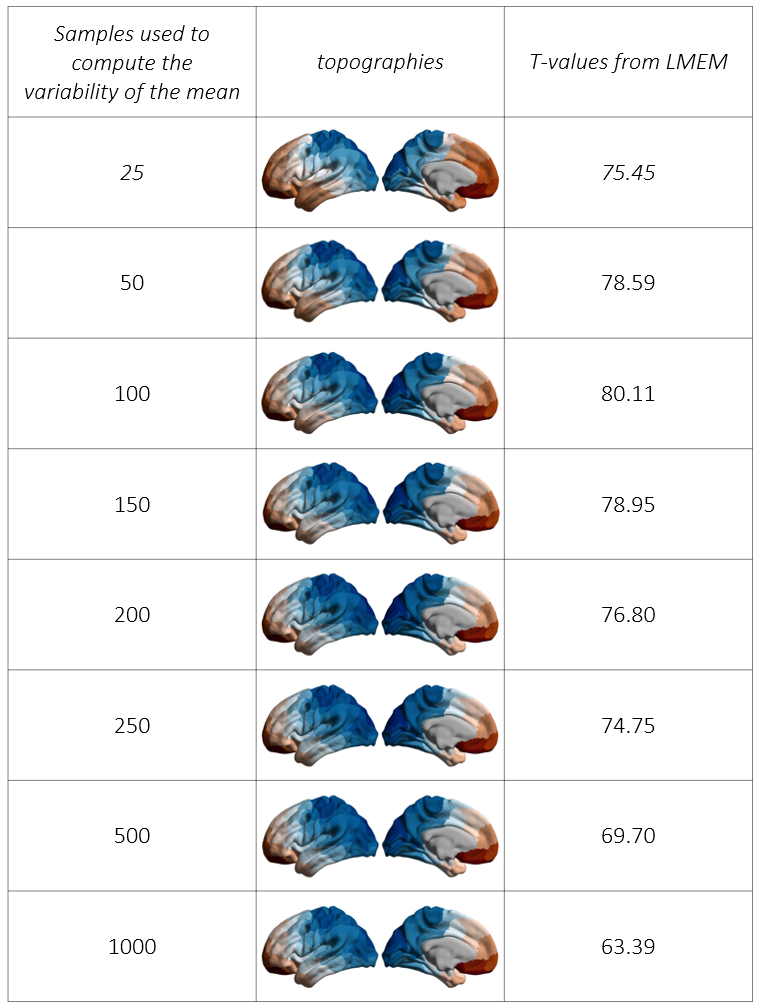

*Figure S3: Group-averaged topographies of the ‘variability of the mean’ feature using different time-window sizes to compute this feature as well as the statistical results for the main effect of archetypal cortical hierarchy on these features evaluated using LMEM.*

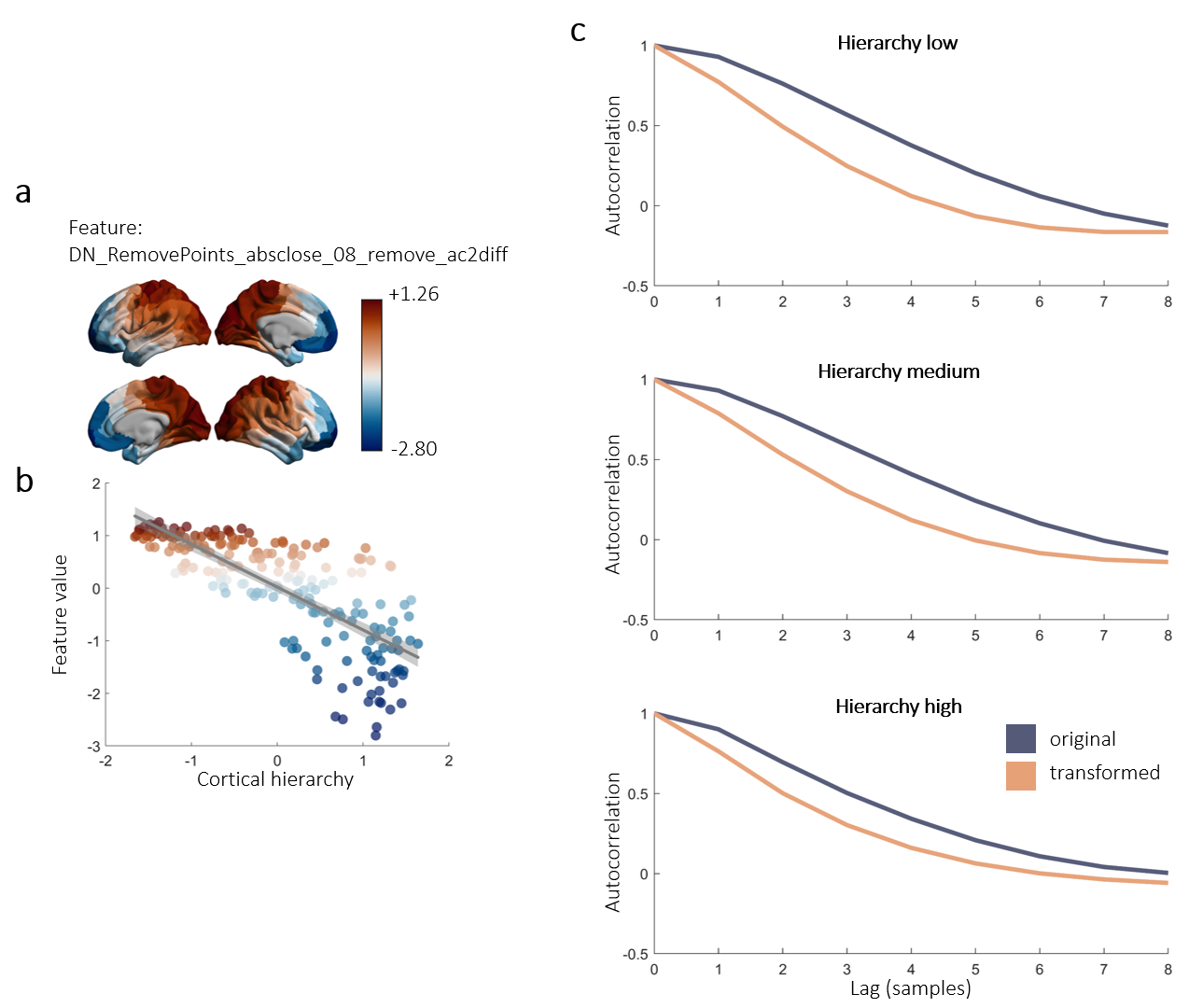

*Figure S4: Visualization of the time-series feature ‘DN_RemovePoints_absclose_08_remove_ac2diff’. From all tested time-series features the presented yielded the highest t-values for a main effect of cortical hierarchy in the LMEM (t = 89.35). The feature describes the difference of the autocorrelation at a lag of 8 samples (27 ms) between the raw time-series and the time-series when 80% of data points closest to the mean are removed. a: Group-averaged z-scored feature plotted across the cortex. Difference in autocorrelation is greatest in motor-related and visual brain areas and decreases towards the prefrontal cortex. b: Group-averaged, z-scored feature plotted across cortical hierarchy. Colors refer to the feature value, therefore red points show high values, while blue indicates low feature values. The gray line describes the linear fit for the displayed data with the 95% confidence bounds as gray shaded area. c: Plotted autocorrelation of the original and the transformed signal across different lags. Data points are averaged across 10 randomly selected participants and across 30 parcels which are lowest, middle, or highest in cortical hierarchy.*

| ***Name*** | ***Estimate*** | ***SE*** | ***tStat*** | ***DF*** | ***pValue*** | ***Lower*** | ***Upper*** |
| --- | --- | --- | --- | --- | --- | --- | --- |
| ***(Intercept)*** | *0.01758707* | *0.0098109* | *1.79260493* | *69996* | *0.07304043* | *-0.00164228* | *0.03681642* |
| ***age*** | *-0.00010162* | *0.00017376* | *-0.58483067* | *69996* | *0.55866347* | *-0.00044219* | *0.00023895* |
| ***hierarchy*** | *-0.68957505* | *0.01051149* | *-65.602012* | *69996* | *0* | *-0.71017755* | *-0.66897255* |
| ***age:hierarchy*** | *0.00398447* | *0.00018617* | *21.4024116* | *69996* | *0* | *0.00361958* | *0.00434936* |

*Table S6: Full statistical results for the LMEM exploring the effects of age, cortical hierarchy, the interaction of both as well as individual random effects on alpha power.*

| ***Name*** | ***Estimate*** | ***SE*** | ***tStat*** | ***DF*** | ***pValue*** | ***Lower*** | ***Upper*** |
| --- | --- | --- | --- | --- | --- | --- | --- |
| ***(Intercept)*** | *3.5794282* | *0.07344352* | *48.7371541* | *69098* | *0* | *3.43547902* | *3.72337738* |
| ***age*** | *-0.02882885* | *0.00125201* | *-23.0261124* | *69098* | *0* | *-0.03128278* | *-0.02637491* |
| ***thick*** | *-1.24836382* | *0.02642591* | *-47.2401528* | *69098* | *0* | *-1.30015855* | *-1.19656909* |
| ***age:thick*** | *0.00925502* | *0.00046124* | *20.0656673* | *69098* | *0* | *0.008351* | *0.01015905* |

*Table S7: Full statistical results for the LMEM exploring the effects of age, cortical thickness, the interaction of both as well as individual random effects on alpha power.*

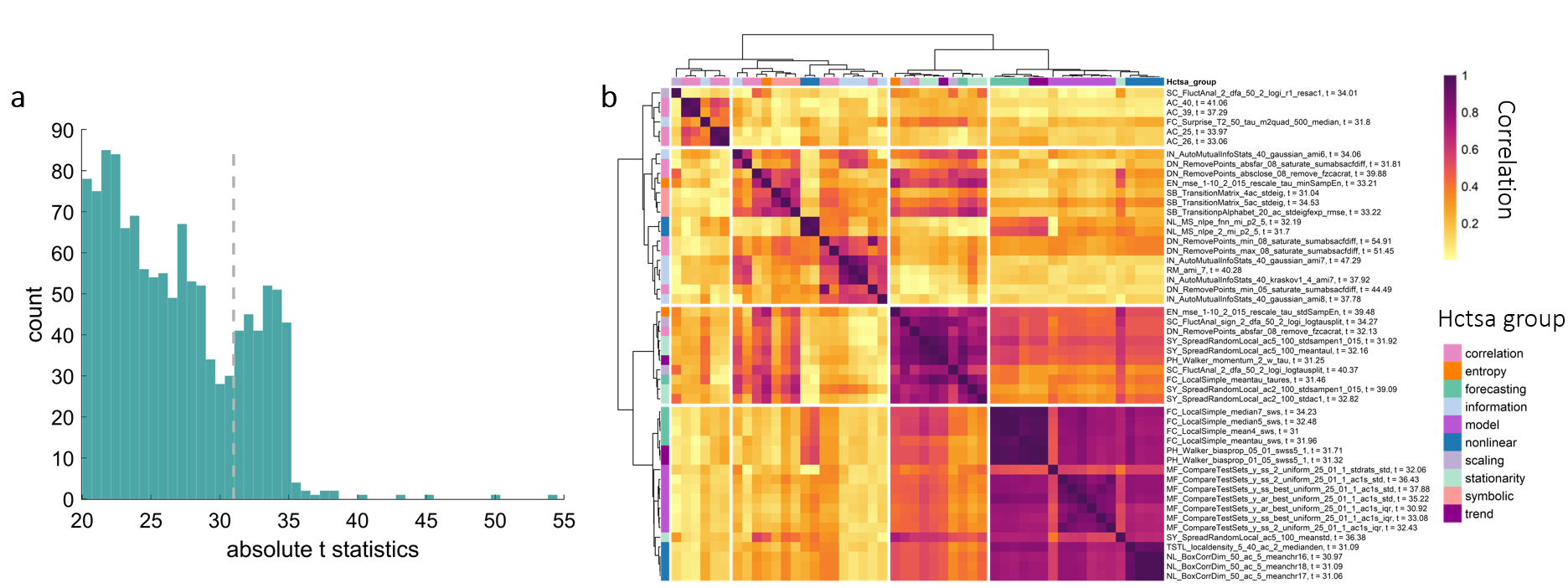

*Figure S5: a: Distribution of the resulting absolute t-values from the LMEM are shown for features which yielded t-values with t>20. The dashed line indicates the threshold for including features in the correlation and cluster analysis.*

*b: Correlation and cluster analysis for the hctsa features with the 50 highest t-values in a LMEM for the interaction effect of the archetypal cortical hierarchy and age. The cluster consists of different subclusters, out of which two features were picked for further analysis.*

| **Feature name** | **t-value LMEM** |
| --- | --- |
| DN_RemovePoints_min_08_saturate_sumabsacfdiff | -54.908 |
| DN_RemovePoints_max_08_saturate_sumabsacfdiff | -51.453 |
| IN_AutoMutualInfoStats_40_gaussian_ami7 | -47.286 |
| DN_RemovePoints_min_05_saturate_sumabsacfdiff | -44.488 |
| AC_40 | -41.057 |
| SC_FluctAnal_2_dfa_50_2_logi_logtausplit | -40.372 |
| RM_ami_7 | -40.285 |
| DN_RemovePoints_absclose_08_remove_fzcacrat | 39.879 |
| EN_mse_1-10_2_015_rescale_tau_stdSampEn | -39.475 |
| SY_SpreadRandomLocal_ac2_100_stdsampen1_015 | 39.091 |
| IN_AutoMutualInfoStats_40_kraskov1_4_ami7 | -37.92 |
| MF_CompareTestSets_y_ss_best_uniform_25_01_1_ac1s_std | 37.878 |
| IN_AutoMutualInfoStats_40_gaussian_ami8 | -37.778 |
| AC_39 | -37.288 |
| MF_CompareTestSets_y_ss_2_uniform_25_01_1_ac1s_std | 36.428 |
| SY_SpreadRandomLocal_ac5_100_meanstd | -36.384 |
| MF_CompareTestSets_y_ar_best_uniform_25_01_1_ac1s_std | 35.221 |
| SB_TransitionMatrix_5ac_stdeig | -34.529 |
| SC_FluctAnal_sign_2_dfa_50_2_logi_logtausplit | -34.267 |
| FC_LocalSimple_median7_sws | 34.229 |
| IN_AutoMutualInfoStats_40_gaussian_ami6 | -34.055 |
| SC_FluctAnal_2_dfa_50_2_logi_r1_resac1 | -34.012 |
| AC_25 | 33.973 |
| SB_TransitionpAlphabet_20_ac_stdeigfexp_rmse | -33.215 |
| EN_mse_1-10_2_015_rescale_tau_minSampEn | 33.209 |
| MF_CompareTestSets_y_ss_best_uniform_25_01_1_ac1s_iqr | 33.084 |
| AC_26 | 33.062 |
| SY_SpreadRandomLocal_ac2_100_stdac1 | 32.817 |
| FC_LocalSimple_median5_sws | 32.482 |
| MF_CompareTestSets_y_ss_2_uniform_25_01_1_ac1s_iqr | 32.432 |
| NL_MS_nlpe_fnn_mi_p2_5 | -32.19 |
| SY_SpreadRandomLocal_ac5_100_meantaul | -32.155 |
| DN_RemovePoints_absfar_08_remove_fzcacrat | 32.134 |
| MF_CompareTestSets_y_ss_2_uniform_25_01_1_stdrats_std | 32.058 |
| FC_LocalSimple_meantau_sws | 31.957 |
| SY_SpreadRandomLocal_ac5_100_stdsampen1_015 | 31.918 |
| DN_RemovePoints_absfar_08_saturate_sumabsacfdiff | -31.808 |
| FC_Surprise_T2_50_tau_m2quad_500_median | -31.799 |
| PH_Walker_biasprop_05_01_swss5_1 | 31.712 |
| NL_MS_nlpe_2_mi_p2_5 | -31.702 |
| FC_LocalSimple_meantau_taures | -31.463 |
| PH_Walker_biasprop_01_05_swss5_1 | 31.325 |
| PH_Walker_momentum_2_w_tau | -31.25 |
| NL_BoxCorrDim_50_ac_5_meanchr18 | -31.087 |
| TSTL_localdensity_5_40_ac_2_medianden | -31.085 |
| NL_BoxCorrDim_50_ac_5_meanchr17 | -31.055 |
| SB_TransitionMatrix_4ac_stdeig | -31.038 |
| FC_LocalSimple_mean4_sws | 31 |
| NL_BoxCorrDim_50_ac_5_meanchr16 | -30.97 |
| MF_CompareTestSets_y_ar_best_uniform_25_01_1_ac1s_iqr | 30.917 |

*Table S8: List of the 50 hctsa time-series features with the highest absolute t-values in a LMEM for an interaction effect of the generalized archetypal cortical hierarchy map and age. Names and t-values are listed, ranked by absolute t-value.*

| ***Name*** | ***Estimate*** | ***SE*** | ***tStat*** | ***DF*** | ***pValue*** | ***Lower*** | ***Upper*** |
| --- | --- | --- | --- | --- | --- | --- | --- |
| ***(Intercept)*** | *-0.01606582* | *0.01070665* | *-1.50054644* | *69996* | *0.13347742* | *-0.03705083* | *0.00491919* |
| ***age*** | *0.00021274* | *0.00018963* | *1.12188841* | *69996* | *0.26191374* | *-0.00015893* | *0.0005844* |
| ***hierarchy*** | *0.62992812* | *0.0114712* | *54.9138653* | *69996* | *0* | *0.60744459* | *0.65241166* |
| ***age:hierarchy*** | *-0.00834132* | *0.00020317* | *-41.0565292* | *69996* | *0* | *-0.00873952* | *-0.00794311* |

*TableS9: Full statistical results for the LMEM exploring the effects of age, cortical hierarchy, the interaction of both as well as individual random effects on autocorrelation at a lag of 133 ms.*

| ***Name*** | ***Estimate*** | ***SE*** | ***tStat*** | ***DF*** | ***pValue*** | ***Lower*** | ***Upper*** |
| --- | --- | --- | --- | --- | --- | --- | --- |
| ***(Intercept)*** | *-2.65765827* | *0.07570203* | *-35.1068286* | *69098* | *0* | *-2.80603412* | *-2.50928241* |
| ***thick*** | *0.93719652* | *0.02723855* | *34.4069944* | *69098* | *0* | *0.88380901* | *0.99058402* |
| ***age*** | *0.02738976* | *0.00129051* | *21.2240187* | *69098* | *0* | *0.02486037* | *0.02991916* |
| ***thick:age*** | *-0.00932687* | *0.00047542* | *-19.6181392* | *69098* | *0* | *-0.01025869* | *-0.00839504* |

*TableS10: Full statistical results for the LMEM exploring the effects of age, cortical thickness, the interaction of both as well as individual random effects on autocorrelation at a lag of 133 ms.*

| ***Name*** | ***Estimate*** | ***SE*** | ***tStat*** | ***DF*** | ***pValue*** | ***Lower*** | ***Upper*** |
| --- | --- | --- | --- | --- | --- | --- | --- |
| ***(Intercept)*** | *0.01140663* | *0.00085586* | *13.3277448* | *69996* | *0* | *0.00972915* | *0.01308411* |
| ***age*** | *-8.2451E-06* | *1.5158E-05* | *-0.54393934* | *69996* | *0.58648495* | *-3.7955E-05* | *2.1465E-05* |
| ***hierarchy*** | *-0.0047407* | *0.00010107* | *-46.9059198* | *69996* | *0* | *-0.00493879* | *-0.0045426* |
| ***age:hierarchy*** | *3.2463E-05* | *1.79E-06* | *18.1353593* | *69996* | *0* | *2.8954E-05* | *3.5971E-05* |

*TableS11: Full statistical results for the LMEM exploring the effects of age, cortical hierarchy, the interaction of both as well as individual random effects on auto mutual information at a lag of 133 ms.*

| ***Name*** | ***Estimate*** | ***SE*** | ***tStat*** | ***DF*** | ***pValue*** | ***Lower*** | ***Upper*** |
| --- | --- | --- | --- | --- | --- | --- | --- |
| ***(Intercept)*** | *0.04047999* | *0.00111473* | *36.3137401* | *69098* | *0* | *0.03829512* | *0.04266486* |
| ***thick*** | *-0.01022341* | *0.0002554* | *-40.0296543* | *69098* | *0* | *-0.01072398* | *-0.00972283* |
| ***age*** | *-0.00027844* | *1.9508E-05* | *-14.273022* | *69098* | *0* | *-0.00031668* | *-0.00024021* |
| ***thick:age*** | *8.9704E-05* | *4.4809E-06* | *20.0192638* | *69098* | *0* | *8.0921E-05* | *9.8486E-05* |

*TableS12: Full statistical results for the LMEM exploring the effects of age, cortical thickness, the interaction of both as well as individual random effects on auto mutual information at a lag of 133 ms.*

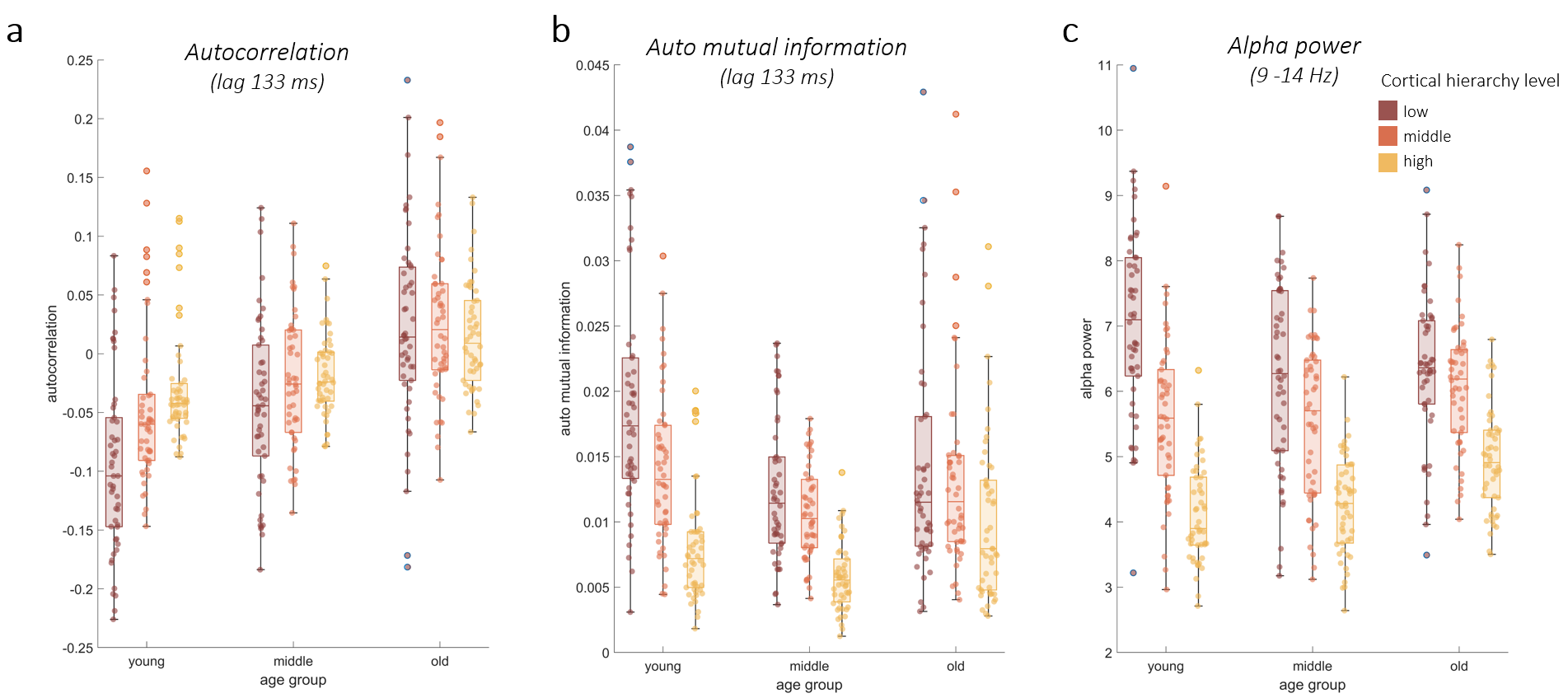

*Figure S6: Visualization of the interaction of the archetypal cortical hierarchy and age on a: autocorrelation (lag 133 ms), b: auto mutual information (lag 133 ms), c: alpha power (8-13 Hz). In each panel the feature value is group-averaged across age groups (18-28, 49-58, 79-88 years old) and parcel hierarchy levels (averaged across ten parcels which are lowest, middle, or highest in cortical hierarchy).*

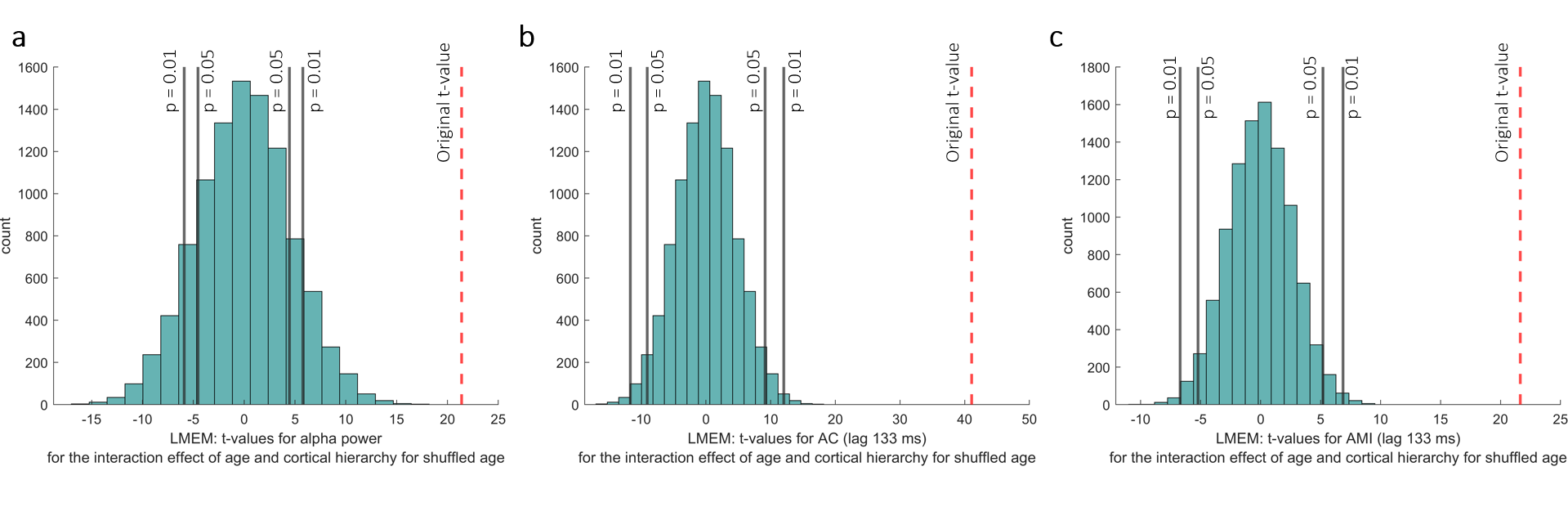

*Figure S7: Histograms of t-values as results of LMEMs for the interaction effect of cortical hierarchy and age on features when ages were shuffled randomly 10.000 times. This way the relationship between the age and cortical hierarchy was removed. Gray lines denote the significance level of ɑ = 5% and ɑ = 1%. For all three features the original t-value indicates a statistical significance of p << 0.001.*
